# Motor adaptation is reduced by symbolic compared to sensory feedback

**DOI:** 10.1101/2024.06.28.601293

**Authors:** Yifei Chen, Sabrina J. Abram, Richard B. Ivry, Jonathan S. Tsay

**Affiliations:** Department of Psychology, University of California, Berkeley; Helen Wills Neuroscience Institute, University of California, Berkeley; Department of Psychology, Carnegie Mellon University

## Abstract

Motor adaptation – the process of reducing motor errors through feedback and practice – is an essential feature of human competence, allowing us to move accurately in dynamic and novel environments. Adaptation typically results from sensory feedback, with most learning driven by visual and proprioceptive feedback that arises with the movement. In humans, motor adaptation can also be driven by symbolic feedback. In the present study, we examine how implicit and explicit components of motor adaptation are modulated by symbolic feedback. We conducted three reaching experiments involving over 400 human participants to compare sensory and symbolic feedback using a task in which both types of learning processes could be operative (Experiment 1) or tasks in which learning was expected to be limited to only an explicit process (Experiments 2 and 3). Adaptation with symbolic feedback was dominated by explicit strategy use, with minimal evidence of implicit recalibration. Even when matched in terms of information content, adaptation to rotational and mirror reversal perturbations was slower in response to symbolic feedback compared to sensory feedback. Our results suggest that the abstract and indirect nature of symbolic feedback disrupts strategic reasoning and/or refinement, deepening our understanding of how feedback type influences the mechanisms of sensorimotor learning.

## Introduction

Motor adaptation – the process of reducing motor errors through feedback and practice – enables us to flexibly move in dynamic and novel environments (Krakauer et al., 2019; Shadmehr et al., 2010; Torres-Oviedo et al., 2011). For example, motor adaptation enables a golfer to adjust her swing to accommodate changes in terrain and wind conditions, and similarly, allows a marathon runner to maintain consistent force output despite increasing muscle fatigue.

Motor adaptation is not a singular process but instead, entails the operation of multiple learning processes. Paralleling the memory literature, one broad distinction can be made between processes that are under conscious control and those that operate outside awareness: Whereas explicit strategy use can allow the agent to reduce performance errors in a volitional and conscious manner (Benson et al., 2011; Hegele & Heuer, 2010; Taylor et al., 2014), implicit recalibration keeps our movements finely calibrated in an automatic and subconscious manner. Indeed, the interplay of explicit and implicit processes in motor adaptation has been the focus of many studies over the past decade (Tsay et al., 2023).

Implicit and explicit learning processes exhibit distinct properties. Whereas implicit recalibration is a relatively rigid process, capable of producing limited, incremental changes in behavior, explicit strategy use is remarkably flexible, capable of producing rapid changes that can be quite dramatic (Bond & Taylor, 2015; Huberdeau et al., 2015). Moreover, while implicit recalibration is highly sensitive to the timing of the feedback and is severely attenuated when the feedback is delayed, strategy-based learning is minimally impacted by manipulations of the timing of the feedback (Brudner et al., 2016; Hadjiosif et al., 2023; Hinder et al., 2008; Honda et al., 2012; Kitazawa et al., 1995; Tsay, Schuck, et al., 2022; Wang, Avraham, et al., 2024).

Learning in most contexts is based on visual and proprioceptive feedback that arises during the movement. These forms of sensory feedback convey motor errors through direct sensory experience: The archer sees their shot off to the left of the bullseye or the guitarist feels their fingers misplaced for the desired chord. Learning, at least for humans, can also be driven by symbolic feedback in which the feedback is conveyed in an indirect, abstract manner. For example, the golfer taking a short cut over the trees might hear a collective groan from the crowd and be fearful that their shot has landed in the pond just in front of the green. Or a blindfolded dart thrower when aiming for the “15”, would know they were too high when informed that the dart was in the “13” slot. While sensory feedback can be exploited by both implicit and explicit adaptation processes (Kim et al., 2018; Neville & Cressman, 2018; Tsay et al., 2021; Tsay, Kim, Haith, et al., 2022), it is unclear if the same holds for symbolic feedback.

In this study, we posed two questions: First, what learning processes are elicited by symbolic feedback? This question remains unresolved because previous studies using symbolic feedback have not used tasks designed to dissociate implicit and explicit learning processes (Galea et al., 2015; Izawa & Shadmehr, 2011; Larssen et al., 2022; Nikooyan & Ahmed, 2015; Therrien et al., 2016; Uehara et al., 2019; Yin et al., 2023). For example, previous studies did not include manipulations such as asking participants to verbally report where they aimed to measure strategy use (Taylor et al., 2014), or instruct participants to forgo strategy use and reach directly to the target when measuring the aftereffect once the perturbation was removed (Maresch et al., 2020; Werner et al., 2015). The only study that has employed such methods found that symbolic feedback was dominated by explicit strategy use (Butcher & Taylor, 2018). In this manuscript, we sought to verify this finding with a substantially larger sample size and with experimental manipulations that provide a more detailed characterization of adaptation driven by symbolic feedback.

Second, is the efficiency of the adaptation process elicited by symbolic feedback comparable to that elicited by sensory feedback? This question has not been clearly answered, given that previous studies have not matched the information content conveyed between sensory and symbolic feedback: Whereas sensory feedback has conveyed both error direction and magnitude information, symbolic feedback has been limited to either magnitude or direction (Butcher & Taylor, 2018; Nikooyan & Ahmed, 2015).

To address these questions, we conducted three motor adaptation experiments involving over 400 human participants. In Experiment 1 we manipulated the size of the rotational perturbation (30°, 60°, 90°), with movement feedback conveyed symbolically via a numerical score. If symbolic feedback elicits implicit recalibration, we expect to observe large and robust aftereffects across all three perturbation sizes, a persistent change in hand angle away from the target after the perturbation is removed and participants are instructed to reach directly towards to the target. Moreover, similar to what is observed with sensory feedback, the magnitude of this aftereffect should be similar for all three perturbation sizes. Conversely, if symbolic feedback is dominated by explicit strategy, we expect learning to scale with perturbation sizes but result in minimal aftereffects. In Experiments 2 and 3, we examined whether the indirect and abstract nature of symbolic feedback, compared to direct and concrete nature of sensory feedback, impacts the discovery of a successful explicit strategy. To this end, we delayed the presentation of feedback to isolate explicit strategy, allowing us to directly contrast learning in response to sensory and symbolic feedback while tightly matching the error information conveyed (i.e., magnitude and direction).

## Methods

### Participants and apparatus

A total of 415 participants completed the study (Female: 216; Male: 174; Other: 25; 24.74 ± 0.17 years old). Participants were recruited on a web-based crowdsourcing platform (www.prolific.com) and were compensated at $12.00/hour. We limited recruitment to participants who 1) have a minimum 95% approval ratings and 2) spoke English as their first language. While handedness was not a recruitment criterion (Right-handed: 362; Left-handed: 46; Ambidextrous: 9), our key findings remained robust even when the analyses were limited to right-handed participants (see Supplemental Section: Figure S1).

The sample size in each experiment were informed by similar web-based motor adaptation studies (Avraham et al., 2021; Wang, Avraham, et al., 2024; Warburton et al., 2023), as well as considerations for counterbalancing. Note that our sample size is significantly greater than comparable in-lab motor adaptation studies (∼40 participants) (Butcher & Taylor, 2018; Larssen et al., 2022; Nikooyan & Ahmed, 2015).

The experiment was created using the OnPoint platform, a package for running customized online motor learning experiments with JavaScript. Participants completed the web-based experiment via an internet browser with their own devices (Trackpad: 332; Optical mouse: 83; Trackball: 2). Our past online studies have shown that neither the type of browser nor the type of pointing device significantly impacts performance (Tsay et al., 2024). The size and position of the visual stimuli were dependent on the individual’s monitor size. For ease of interpretation, all stimulus parameters detailed below were based on an average 13-inch computer monitor.

### General procedure

For each trial, participants were asked to position their cursor, a white dot (diameter = 0.4 cm), inside a white starting ring (diameter = 0.5 cm) which was located at the center of the screen. Once the cursor was moved inside, the starting ring was filled. After holding the cursor inside the starting ring for 500 ms, the cursor was blanked, eliminating visual feedback, and a blue circular target (diameter = 0.4 cm) appeared along an invisible ring with a radius of 8 cm relative to the starting ring. The blue target could appear in one of the three target locations on the invisible ring. The sequence of target locations was presented pseudo-randomly within each movement cycle (i.e., 1 movement cycle = 3 reaches: 1 reach to each target location).

### Feedback

Feedback was provided immediately (Experiment 1) or 800 ms after movement termination (Experiments 2-3) and remained visible for 1 s. Symbolic feedback was provided via a numerical score. How the numerical score was calculated varied between experiments: In Experiment 1, the score conveyed only error magnitude, the absolute distance between the participant’s movement endpoint and the target, with the score rounded to the nearest integer. To make the scores easier for the participants to understand, we normalized the range of scores from a minimum of 0 points (hand angle 180° away from the target) to a maximum of 100 points (hand angle at the bullseye of the target). In Experiments 2 and 3, symbolic feedback conveyed both error magnitude and direction. Here the score could range from –180° to 179°. As such, 0 was the best possible score in these experiments, with negative values indicating a counterclockwise error and positive values a clockwise error (e.g., a score of +60 signified that the endpoint hand angle was 60° clockwise from the optimal location).

We also included sensory feedback conditions in Experiments 2 and 3. Sensory feedback was provided via a white cursor that appeared on the ring, indicating the participant’s hand position when the movement amplitude was 8 cm (veridical feedback trials) or displaced from that hand position by the visual perturbation (rotation trials).

### Experiment 1

Participants (N = 184; Female: 93; Male: 77; Other: 14; 25.06 ± 0.29 years old) were randomly assigned to one of the three perturbation groups (30° rotation: 77; 60° rotation: 51; 90° rotation: 56). Two participants whose average hand angles were 5 standard deviations away from the group means were excluded. We counterbalanced the direction of the perturbation (clockwise or counterclockwise) across participants within each perturbation group. There were three target locations: 30° (upper-right quadrant), 150° (upper-left quadrant), and 270° (straight down).

Movement feedback was always provided symbolically in the form of a numerical score that ranged between 0 and 100 points. There were three blocks: baseline veridical feedback (30 trials; 10 cycles), rotated feedback (150 trials; 50-cycles), and no-feedback aftereffect (30 trials; 10 cycles). In the baseline block, participants were familiarized with the basic reaching procedure and the symbolic feedback. Participants were provided the following instructions: “Move directly to the target. You will be rewarded based on your accuracy (max score = 100 points).” In the perturbation block, the score was based on the rotated (30°, 60° or 90°) endpoint hand position. Thus, to get 100 points on a trial, a participant in the 60° clockwise perturbation group would have to move 60° counterclockwise to the target. Participants were instructed: “Move somewhere away from the target. Find the movement direction that yields 100 points.” In the aftereffect block, the perturbation was removed. Participants were given the following instructions: “Move directly to the blue target, and do not aim away from the target.” There was no feedback presented during the aftereffect block.

### Experiment 2

Participants (N = 110; Female: 58; Male: 46; Other: 6; 24.33 ± 0.31 years old) were randomly assigned to one of the two groups that differed in terms of feedback type (Symbolic: 53; Sensory: 57). For both groups, the feedback provided vectorial information (magnitude and direction). Importantly, the feedback was presented 800 ms after the hand had reached the target amplitude – a manipulation that greatly attenuates or even eliminates implicit recalibration (Brudner et al., 2016; Tsay, Schuck, et al., 2022). In this way, we sought to focus on comparing explicit strategy use in response to sensory or symbolic feedback. We counterbalanced the direction of the perturbation (clockwise or counterclockwise) across participants within each feedback group. The target locations were the same as Experiment 1.

There were three experimental blocks: Baseline veridical feedback (30 trials; 10 cycles), delayed 60° rotated feedback (150 trials; 50 cycles), and no-feedback aftereffect (30 trials; 10 cycles). Unlike Experiment 1, we introduced 12 instruction familiarization trials at the beginning of the baseline block to ensure that participants fully understood the error information conveyed by the feedback. During the first six familiarization trials, the following instructions accompanied the feedback display: “You missed the target by 10° in the clockwise direction” (Sensory group) or “You missed the target by 10° in the clockwise direction; a positive score signifies a clockwise error, and a negative score signifies a counterclockwise error” (Symbolic group). Participants advanced to the next trial by pressing the space bar. In the final six familiarization trials, participants were required to report the direction of their error after the feedback was presented (press ‘a’ for a clockwise error; press ‘b’ for a counterclockwise error). The experiment was terminated if participants provided inaccurate responses for more than two of these six trials.

### Experiment 3

Participants (N = 121; Female: 65; Male: 51; Other: 5; 24.62 ± 0.30 years old) were assigned to one of the two groups receiving symbolic (50) or sensory (71) feedback. The feedback was provided in the same manner as Experiment 2. We designed Experiment 3 to contrast learning performance in response to symbolic and sensory feedback conveying a mirror reversal perturbation. Specifically, the endpoint hand position was mirror-reversed across either the horizontal or vertical axis (reversal axes counterbalanced among participants). For example, in the horizontal mirror condition, if participants reached to the 30° target, their endpoint hand position would be reflected to the 330° location, resulting in a 60° error. Similarly, in the vertical mirror condition, if participants reached to the 240° target, their endpoint hand position will be reflected to the 300° location, also resulting in a 60° error. Note that, unlike a rotational perturbation, re-aiming in the opposite direction of the error (sensory of symbolic) would increase the error in response to the mirror transformation.

The target locations were dependent on the mirror reversal axis to maintain a consistent 60° error if participants move directly to the target (Figure 3a, b, right panel). Specifically, participants experiencing a horizontal mirror reversal moved to targets located at 30°, 150°, and 210°, or to targets located at 30°, 150°, and 330° (counterbalanced across participants). Likewise, participants experiencing a vertical mirror reversal moved to targets located at 60°, 120°, and 240°, or to targets located at 60°, 120°, and 300° (counterbalanced across participants).

There were three blocks: Baseline veridical feedback (30 trials; 10 cycles), delayed mirror-reversed feedback (150 trials; 50 cycles), and no-feedback aftereffect (30 trials; 10 cycles). To ensure participants fully understood the feedback provided, we incorporated an instruction familiarization block similar to that of Experiment 2.

### Data analysis

We focused our analyses on the hand position data recorded when the movement amplitude reached the target radius. These data used to calculate our main dependent variable, hand angle, the difference between the hand position and the target. Hand angles across different perturbation directions were flipped such that positive hand angles always signified changes in heading angle that nullifies the perturbation. Hand angles were also baseline subtracted to correct for small idiosyncratic movement biases (Vindras et al., 1998; Wang, Morehead, et al., 2024). Baseline performance included all 10 cycles of the baseline veridical feedback block (trials 1 – 30). Early adaptation was operationally defined as the first 10 cycles of the perturbation block (trials 31 – 60) and late adaptation was defined as the last 10 cycles of the perturbation block (trials 151 – 180). Aftereffect performance included all 10 cycles of the aftereffect block (trials 181 – 210).

We also used a continuous performance measure to compare the groups, implementing a cluster-based permutation test on the hand angle and reaction time data (Breska & Ivry, 2019; Sassenhagen & Draschkow, 2019; Tsay et al., 2020). The test consisted of two steps. First, a F-test (comparing >2 experimental conditions) or t-test (comparing 2 experimental conditions) was performed for each movement cycle across experimental conditions to identify clusters showing a significant difference. Clusters were defined as epochs in which the p-value from the F– or t-tests were less than 0.05 for at least two consecutive cycles. The F or t values were then summed up across cycles within each cluster, yielding a combined cluster score. Second, to assess the probability of obtaining a cluster of consecutive cycles with significant p-values, we performed a permutation test. Specifically, we generated 1000 permutations by shuffling the condition labels. For each shuffled permutation, we calculated the sum of the F– or t-scores. Doing this for 1000 permutations generated a distribution of scores. The proportion of random permutations which resulted in a F-score or a t-score that was greater than or equal to that obtained from the data could be directly interpreted as the p-value. Clusters with *p_perm_* < 0.05 are reported. We also reported the minimum effect size of all clusters (Cohen’s d for between-participant comparisons; and 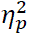 for main effects).

We performed a subgroup analysis exclusively on “learners” (Brudner et al., 2016; Jang et al., 2023; Tsay, Schuck, et al., 2022). Using a pair of liberal criteria, learners were defined as participants whose 1) hand angle was greater than 20% of the perturbation (e.g., greater than 12° of the 60° perturbation) and 2) demonstrated a significant change in hand angle in the direction that correctly counteracts the perturbation during late adaptation (one tailed paired t-tests between baseline and late adaptation, *t value > 0* and *p < 0.05*). Two participants who constantly aimed toward the opposite direction of the target were manually removed from “learners” by visual inspection. This analysis yielded 112 learners in Experiment 1 (proportion of learners: 61%), 76 learners in Experiment 2 (69%), and 75 learners in Experiment 3 (62%).

### Data and code availability statement

Raw data and analysis code can be openly accessed at: https://osf.io/bpfnh/?view_only=b1f4ba4e5576462c9be266380e2fee8b

## Results

### Experiment 1: Motor adaptation in response to symbolic feedback is dominated by explicit strategy use

We presented unsigned symbolic feedback in Experiment 1, varying the size of the rotational perturbation (30°, 60°, 90°) in a between-group design (Figure 1a). Feedback was limited to a number ranging from 0 (cursor moved in the opposite direction of the target) to 100 (cursor landed on target). We included a no-feedback aftereffect block in which we instructed the participants to reach “directly to the target.” The cardinal signature of implicit recalibration is a residual deviation in hand angle during the aftereffect block, with the magnitude of the effect similar across different perturbation sizes. Signatures of explicit strategy use include 1) the scaling of adaptation across these large perturbation sizes, considering that implicit recalibration should have already saturated by 30° (Bond & Taylor, 2015; Kim et al., 2018) and 2) an immediate and large change in hand angle back towards the target at the start of the aftereffect block.

**Figure 1.**
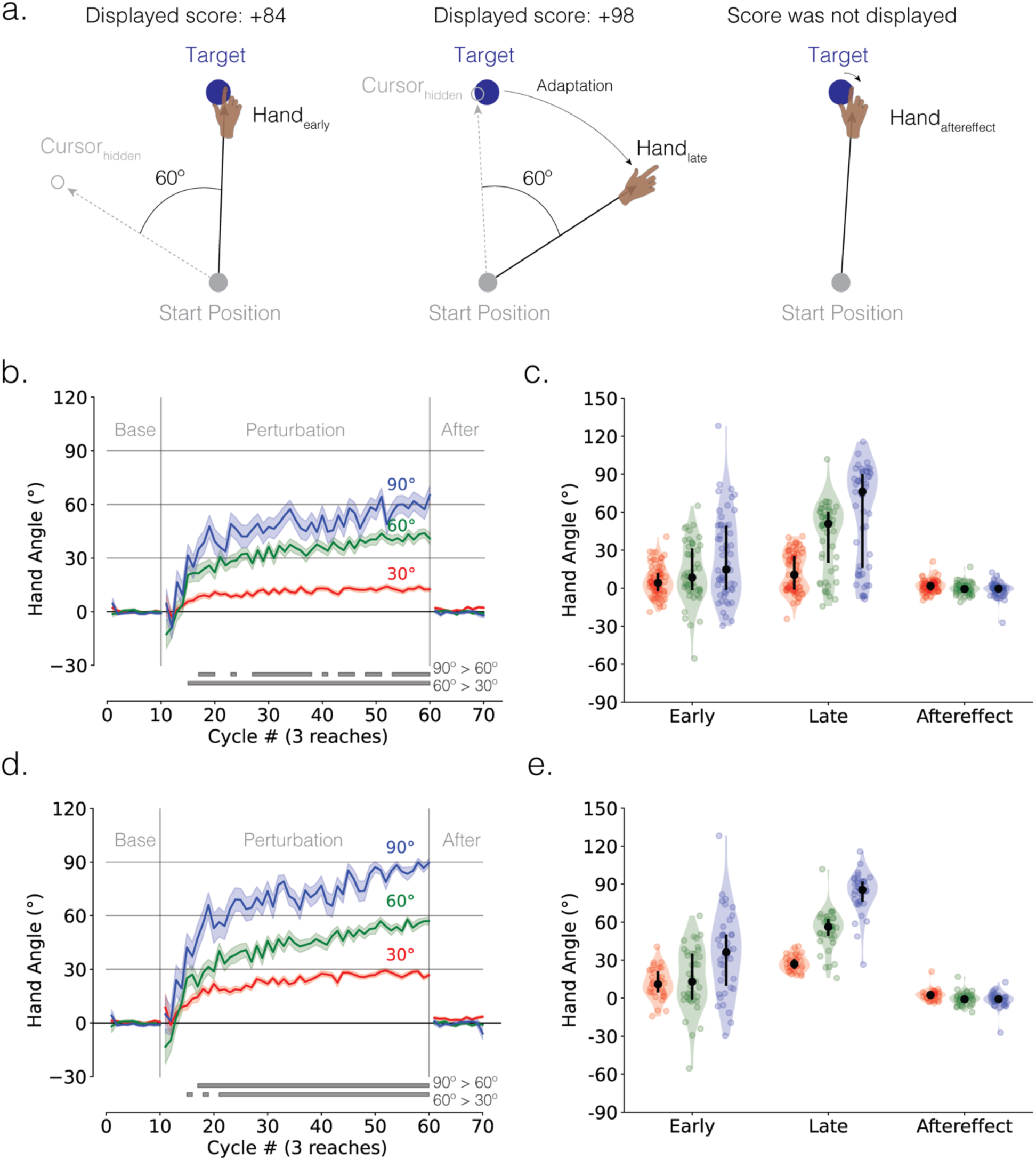
Motor adaptation in response to symbolic feedback is dominated by strategy use. (**a**) Schematic of the web-based rotational perturbation task in Experiment 1. An example of the 60° counterclockwise rotation is provided. Participants are instructed to reach in the direction that maximized points (i.e., 100 points). The left, middle, and right panels display a representative trial from the early adaptation, late adaptation, and aftereffect phases, respectively. **(b)** Mean time courses of hand angle (N = 184; 30°: 77 participants; 60°: 51 participants; 90°: 56 participants). Colors denote different perturbation groups (red = 30°; green = 60°; blue = 90°). Shaded error denoted SEM. Each movement cycle includes three trials (1 reach to each of the three targets). Grey horizontal lines at the bottom indicate clusters showing significant group differences. **(c)** Mean hand angles during early adaptation (i.e., first 10 cycles of the perturbation block), late adaptation (last 10 cycles of the perturbation block), and aftereffect phases (10 cycles of the aftereffect block). Black line denotes median ± IQR. **(d)** Mean time courses and **(e)** mean hand angles of learners (N = 112; 30°: 35 learners; 60°: 39 learners; 90°:38 learners).

Overall, participants improved their scores over the course of the perturbation block, exhibiting a change in hand angle away from the target (Figure 1b). Even though the mean hand angle for each groups fell considerably short of optimal performance during late adaptation, the learning functions scaled with the size of the rotation. Notably, there was a large change in hand angle at the start of the aftereffect block and minimal evidence of a residual aftereffect in all three groups. Together, these results are consistent with the hypothesis that adaptation in response to symbolic feedback is dependent on the use of an explicit re-aiming strategy, with minimal evidence of any implicit recalibration.

These observations were confirmed in a series of statistical tests. In terms of the tests of strategic re-aiming, mean hand angle throughout the perturbation block exhibited a main effect of perturbation size (cluster-based F-test for a main effect of perturbation size, *F*_score_ = 1579.50*, p*_perm_ < 0.05, 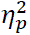 > 0.106). Post-hoc cluster-based t-tests showed that learning was greatest for 90° and smallest for 30° (grey solid lines in Figure 1b: **90° > 60°**: *t*_sum_ > 4.72, *p*_perm_ < 0.05, *d* > 0.2; **60° > 30°**: *t*_sum_ = 302.25, *p*_perm_ < 0.05, *d* = 0.7). We observed similar scaling effects in reaction time, with reaction time being longest for the 90**°** group and shortest for the 30**°** group (Supplemental Figure S2a; **90° > 60°**: *t*_sum_ > 4.13, *p*_perm_ < 0.05, *d* > 0.2; **60° > 30°**: *t*_sum_ > 5.59, *p*_perm_ < 0.05, *d* > 0.2). The scaling of reaction time has been taken to indicate that both the discovery of an explicit strategy becomes more computationally demanding as the perturbation size increases (Guo & Song, 2023; McDougle & Taylor, 2019; Pellizzer & Georgopoulos, 1993). In contrast, varying the perturbation size across the range used in Exp 1 has negligible effects on both the learning functions and reaction time in tasks that isolate implicit recalibration (Marko et al., 2012; Morehead et al., 2017; Tsay et al., 2023).

Further evidence of strategy use is given by the observation that all three groups showed a significant drop in hand angle from late adaptation to aftereffect block (Figure 1c; paired t-tests comparing aftereffect vs late adaptation, **30°**: –10.69 ± 1.69°, *t*(76) = –6.59, *p* < 0.001, *d* = –0.96; **60°**: –41.16 ± 3.82°, *t*(50) = –10.99, *p* < 0.001, *d* = –2.16; **90°**: –60.12 ± 5.22°, *t*(55) = –11.02, *p* < 0.001, *d* = –2.11). Indeed, we observed either minimal or absent aftereffects in the three perturbation groups (Figure 1c; paired t-tests comparing aftereffect vs baseline, **30°**: 1.70 ± 0.55°, *t*(76) = 3.20, *p* = 0.002, *d* = 0.38; **60°**: –0.18 ± 0.64°, *t*(50) = –0.28, *p* = 0.782, *d* = –0.05; **90°**: –1.09 ± 0.74°, *t*(55) = –1.88, *p* = 0.07, *d* = –0.2). Thus, the data suggest that the error information conveyed by symbolic feedback is not sufficient to engage the process underlying implicit recalibration, similar to the findings of Butcher and Taylor (Butcher & Taylor, 2018). Note that the absence of aftereffect is not a peculiarity of conducting remotely experiments over the web, as many web-based studies have elicited robust implicit recalibration in response to a rotational perturbation (Jang et al., 2023; Shyr & Joshi, 2023; Tsay et al., 2024).

As noted above, late adaptation for all groups fell considerable short of optimal performance (paired t-tests comparing late adaptation vs baseline, **30°**: 12.40 ± 1.75°, *t*(76) = 7.33, *p* < .001, *d* = 1.1; **60°**: 40.98 ± 3.78°, *t*(50) = 11.06, *p* < .001, *d* = 2.1; **90°**: 59.02 ± 5.15°, *t*(55) = 10.93, *p* < .001, *d* = 2.1). Inspection of individual data indicated that there were several participants in each group who exhibited minimal improvement, with mean hand angles remaining close to baseline (i.e., towards the target, Figure 1c; **30°**: 55% non-learners; **60°**: 24% non-learners; **90°**: 32% non-learners; See Supplemental Figure 3a for representative non-learners). While the presence of non-learners may point to a general performance issue (e.g., failure to attend to the task), it may also highlight how learning in response to symbolic feedback is unlikely to be automatic and implicit, but instead, explicit and computationally demanding (Fernandez-Ruiz et al., 2011; Huberdeau et al., 2019).

Given that each group appears to be composed of “learners” and “non-learners”, we repeated the key analyses with only the data from the “learners” (see Methods for inclusion criteria). In this restricted, post-hoc analysis, the core observations noted above were even more striking, with late adaptation approaching the full perturbation (paired t-tests against the baseline, **30°**: 27.52 ± 1.04°, *t*(34) = 26.54, *p* < 0.001, *d* = 5.6; **60°**: 53.76 ± 2.43°, *t*(38) = 22.10, *p* < 0.001, *d* = 5.0; **90°**: 83.10 ± 2.59°, *t*(37) = 32.10, *p* < 0.001, *d* = 7.0). The change in hand angle during late adaptation scaled with the size of the perturbation (Figure 1d; **90° > 60°**: *t*_sum_ > 209.98, *p*_perm_ < 0.05, *d* = 0.6; **60° > 30°**: *t*_sum_ > 5.00, *p*_perm_ < 0.05, *d* > 0.3), as did reaction times (Supplemental Figure S2d; **90° > 60°**: *t*_sum_ > 6.98, *p*_perm_ < 0.05, *d* > 0.3; **60° > 30°**: *t*_sum_ > 14.12, *p*_perm_ < 0.05, *d* > 0.3).

Perhaps most interesting, the aftereffects remained negligible in the restricted analysis, being significantly different from zero only in the 30° group (Figure 1e; **30°**: 2.36 ± 0.74°, *t*(34) = 3.22, *p* = 0.003, *d* = 0.5; **60°**: –0.49 ± 0.79°, *t*(38) = –0.62, *p* = 0.536, *d* = –0.1; **90°**: –1.20 ± 0.97°, *t*(37) = –1.70, *p* = 0.100, *d* = –0.3). It is unclear whether this 2° aftereffect in the 30° group is the result of implicit recalibration (van Mastrigt et al., 2023) or a small use-dependent bias caused by repeated movements away from the target (Diedrichsen et al., 2010; Marinovic et al., 2017; Mawase et al., 2017; Tsay, Kim, Saxena, et al., 2022; Wood et al., 2020).

Together, the results of Experiment 1 underscore how motor adaptation in response to symbolic feedback is dominated by, and in most cases limited to, strategy use.

### Experiment 2: Motor adaptation in response to a rotational perturbation is reduced by symbolic compared to sensory feedback

The results of Experiment 1 raise the question of whether the processes underlying strategy discovery differ when triggered by symbolic or sensory feedback. Previous studies comparing sensory and symbolic feedback have not matched the error information (direction and magnitude) conveyed by these different types of feedback (Butcher & Taylor, 2018; Codol et al., 2018; Galea et al., 2015; Holland et al., 2018; Izawa & Shadmehr, 2011; Larssen et al., 2022; Nikooyan & Ahmed, 2015; Therrien et al., 2016; Uehara et al., 2019; Yin et al., 2023). We set out to fill this gap in Experiment 2.

To provide a fair comparison between the two feedback types, we used perturbation conditions that should ensure learning is limited to explicit strategy use. To this end, we used a large perturbation (60° rotation) and presented the feedback 800 ms after the amplitude of the hand movement reached the target distance (Figure 2). The latter manipulation severely attenuates, or even eliminates any contribution of implicit recalibration (Brudner et al., 2016; Kitazawa et al., 1995). Unlike Experiment 1, the symbolic feedback was modified to convey information about both error magnitude and error direction. In this way, we sought to create two conditions that only differed in whether the terminal position of the cursor was indicated by a sensory cue at that position or symbolic feedback indicating that position.

**Figure 2.**
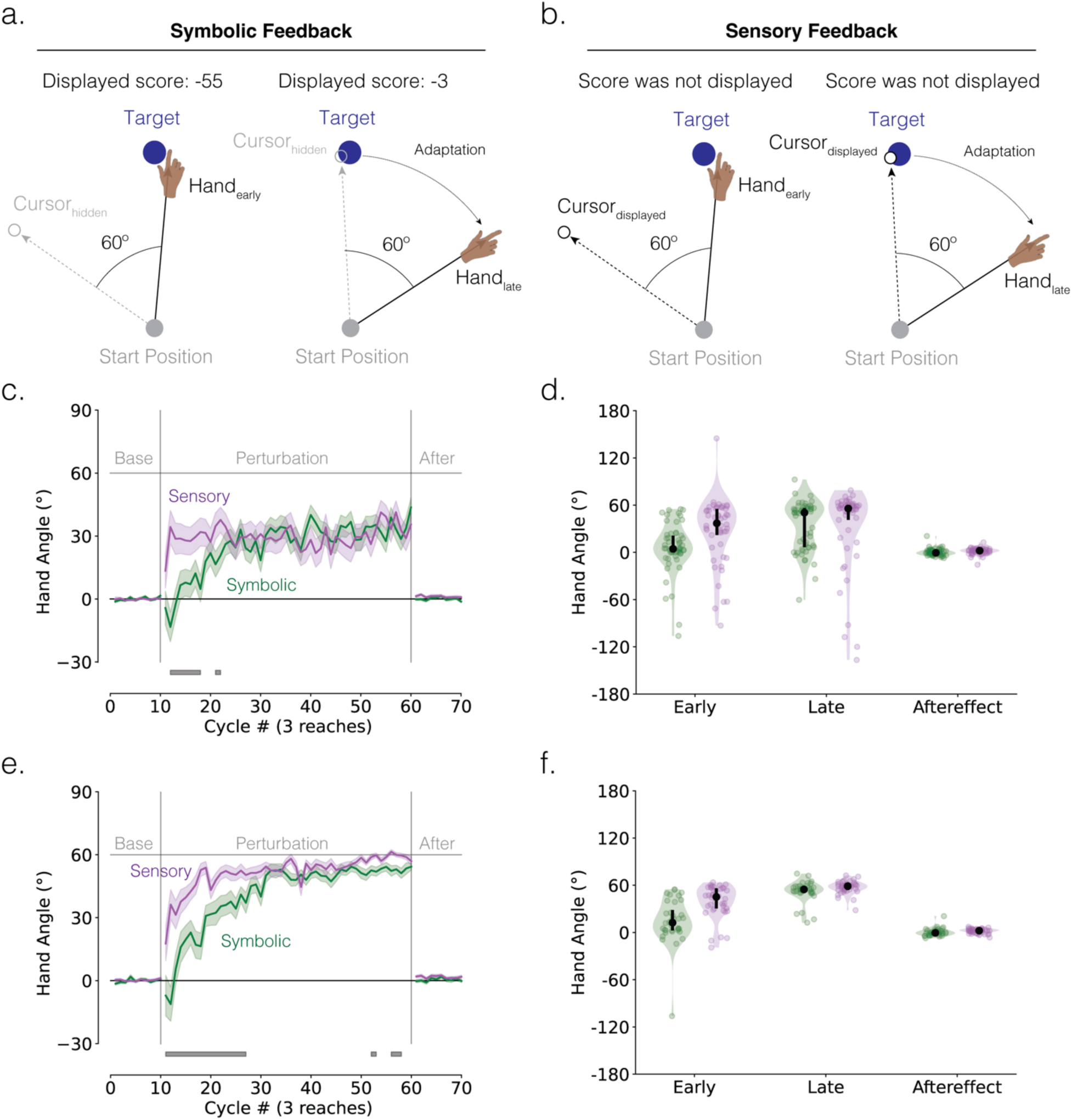
Symbolic feedback reduces explicit strategy use in response to a rotational perturbation. (**a, b**) Schematic of the 60° visuomotor rotation task (Experiment 2). Feedback was delayed to minimize implicit recalibration. Participants were instructed to move in a direction that minimizes error. Error was conveyed via **(a)** symbolic feedback (score magnitude conveys the size of the angular error rounded to the nearest integer; score sign conveys direction, with negative denoting a clockwise error and positive denoting a counterclockwise error) or **(b)** sensory feedback (the magnitude and direction of the error are conveyed by a rotated cursor). Left and right panels denote early and late adaptation, respectively. **(c)** Mean time courses of hand angle (N = 110; Sensory: 53 participants; Symbolic: 57 participants). Colors denote different feedback groups (dark green = symbolic feedback; dark magenta = sensory feedback). Shaded error denoted SEM. Grey horizontal lines at the bottom indicate clusters showing significant group differences. **(d)** Mean hand angles during early adaptation, late adaptation, and aftereffect phases. Black line denotes median ± IQR. **(d)** Mean time courses and mean hand angles of learners (N = 76; Sensory: 40 learners; Symbolic: 36 learners).

The learning functions for the symbolic (dark green) and sensory (dark magenta) groups are shown in Figure 2c. Both groups reached a similar level of asymptotic performance, one that fell short of counteracting the 60° perturbation. However, the sensory group reached this asymptote within just a few movement cycles, a much faster rate than that exhibited by the symbolic group. Both groups exhibited minimal aftereffects when the feedback was removed, confirming that the delayed feedback manipulation was successful in eliminating implicit recalibration.

These observations were verified statistically. First, the cluster-based permutation test demonstrated that the change in hand angle occurred more slowly in response to symbolic feedback compared to sensory feedback, with a significant difference between groups evident in many of the initial perturbation cycles (Figure 2c; cluster-based t-test, *t*_score_ > 4.11, *p*_perm_ < 0.05, *d* > 0.2). This group difference was not significant after cycle 23. Second, both groups exhibited a large drop in hand angle after late adaptation (Figure 2d; paired t-tests comparing aftereffect vs late adaptation, **Sensory**: –32.89 ± 6.84°, *t*(52) = –5.04, *p* < 0.001, *d* = –0.9; **Symbolic**: –34.06 ± 4.14°, *t*(56) = –6.81, *p* < 0.001, *d* = –1.3). Third, neither group exhibited a significant aftereffect (Figure 2d; paired t-tests comparing aftereffect vs baseline, **Sensory**: 1.49 ± 0.56°, *t*(52) = –0.37, *p* = 0.716, *d* = –0.1; **Symbolic**: 0.11 ± 0.58°, *t*(56) = 0.18, *p* = 0.855, *d* = 0.0).

As in Experiment 1, the low asymptotes for both groups were primarily due to fact that some participants in each group failed to come up with the correct strategy by the end of the experiment (**Sensory**: 24% non-learners; **Symbolic**: 37% non-learners). For the “learner” subgroup, late adaptation in both feedback groups approximated the size of the perturbation (Figure 2f; paired t-tests comparing late adaptation vs baseline, **Sensory**: 57.35 ± 1.26°, *t*(39) = 45.53, *p* < 0.001, *d* = 9.8; **Symbolic**: 52.30 ± 2.21°, *t*(35) = 23.62, *p* < 0.001, *d* = 5.5). Importantly, the cluster-based permutation test again demonstrated that learning occurred more slowly in response to symbolic compared to sensory feedback, with a significant hand angle difference between groups evident in both initial and late perturbation cycles (Figure 2e; cluster-based t-test, *t*_score_ > 4.41, *p*_perm_ < 0.05, d > 0.2).

There was a significant drop in mean hand angles at the start of the aftereffect block (Figure 2f: **Sensory**: – 55.58 ± 1.29°, *t*(39) = –43.15, *p* < 0.001, *d* = –9.2; **Symbolic**: –51.93 ± 2.47°, *t*(35) = –20.99, *p* < 0.001, *d* = –5.22), with minimum aftereffect (Figure 2f; paired t-tests comparing aftereffect vs baseline, **Sensory**: 1.78 ± 0.47°, *t*(39) = 3.82, *p* < 0.001, *d* = 0.5; **Symbolic**: 0.36 ± 0.81°, *t*(35) = 0.45, *p* = 0.658, *d* = 0.1).

The results of Experiment 2 show that explicit motor adaptation is slower in response to symbolic compared to sensory feedback even when implicit recalibration is minimized, and the error information is closely matched in terms of direction and magnitude. It appears that the ability to discover a successful re-aiming strategy is hindered when the feedback is presented in an indirect and abstract format.

### Experiment 3: Motor adaptation in response to a mirror reversal perturbation is reduced by symbolic compared to sensory feedback

We designed Experiment 3 to examine if the learning disadvantage observed with symbolic feedback would also be manifest with another type of perturbation. To test this, we used a mirror reversal of the visual feedback, a perturbation known to elicit adaptation mainly through explicit strategy use (Figure 3a-b) (Lillicrap et al., 2013; Telgen et al., 2014; Yang et al., 2021).

**Figure 3.**
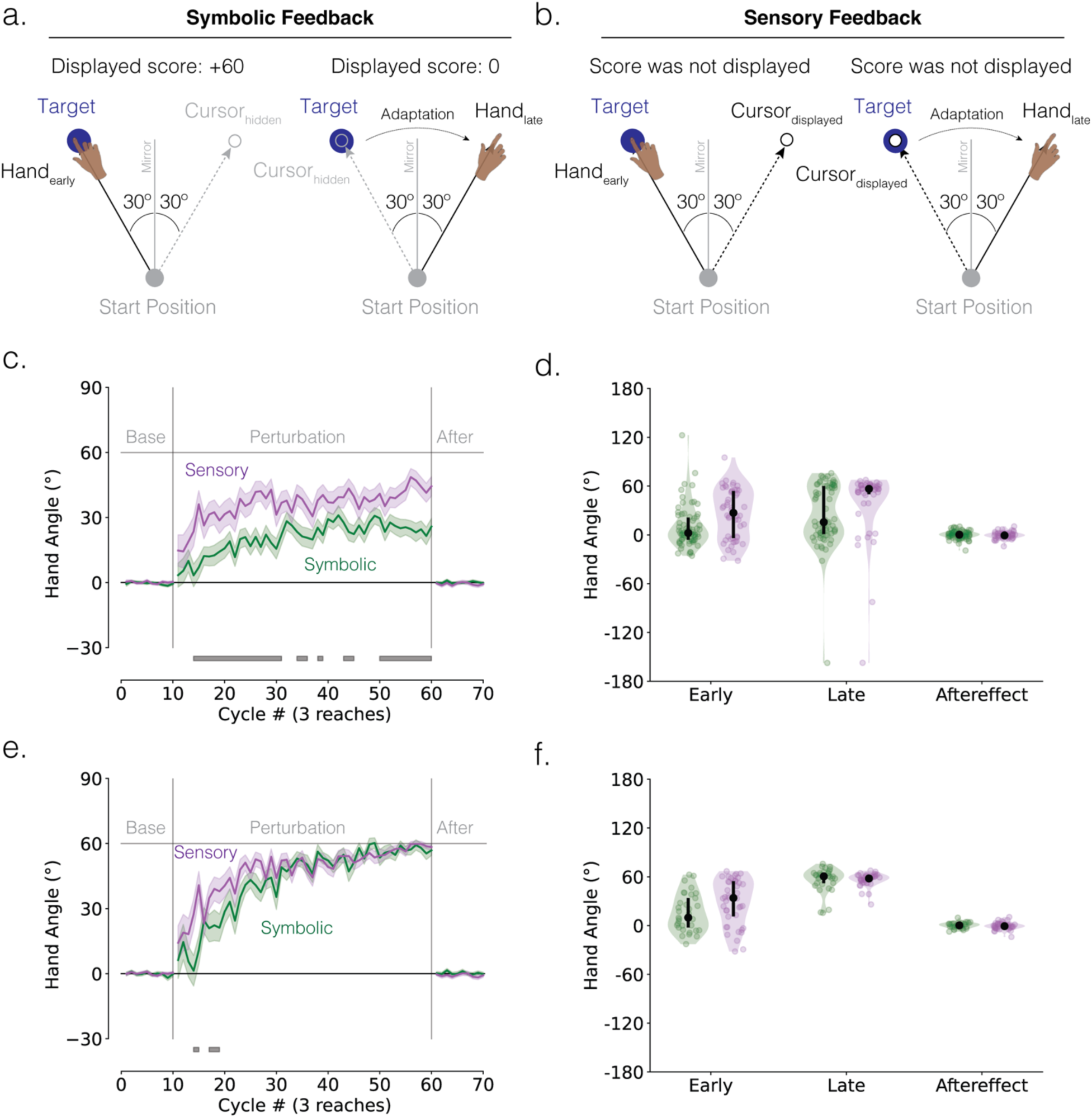
Symbolic feedback reduces explicit strategy use in response to a mirror perturbation. (**a, b**) Schematic of the mirror-reversal perturbation task in Experiment 3. Participants were instructed to move in the direction that minimizes error. Error was conveyed via **(a)** symbolic feedback or **(b)** sensory feedback. Feedback is provided in the same manner as Experiment 2. Left and right panels denote early and late adaptation, respectively. **(c)** Mean time courses of hand angle (N = 121; Sensory: 50 participants; Symbolic: 71 participants). Colors denote different feedback groups (dark green = symbolic feedback; dark magenta = sensory feedback). Shaded error denoted SEM. Grey horizontal lines at the bottom indicate clusters showing significant group differences. **(d)** Mean hand angles during early adaptation, late adaptation, and aftereffect phases. Black line denotes median ± IQR. **(d)** Mean time courses and **(e)** mean hand angles of learners (N = 75; Sensory: 42 learners; Symbolic: 33 learners).

When we compared the learning functions between the symbolic feedback and sensory feedback groups, our key results mirrored (pun intended) those of Experiment 2 (Figure 3c). The cluster-based permutation test revealed that the change in hand angle occurred slower in response to symbolic feedback compared to sensory feedback, with a significant difference between groups evident throughout the entire perturbation block (Figure 3c; cluster-based t-test, all participants: *t*_score_ > 4.94, *p_perm_* < 0.05, *d* > 0.2). Consistent with the assumption that performance on this task is strategy driven, both groups exhibited a significant drop in hand angle from the late adaptation to aftereffect phases (Figure 3d; paired t-tests against baseline: **Sensory**: – 42.71 ± 5.74°, *t*(49) = –6.30, *p* < 0.001, *d* = –1.3; **Symbolic**: –24.63 ± 4.47°, *t*(70) = –5.26, *p* < 0.001, *d* = – 0.9). Neither group showed a significant aftereffect (Figure 3d; paired t-tests against baseline: **Sensory**: – 0.45 ± 0.58°, *t*(49) = –0.78, *p* = 0.441, *d* = –0.1; **Symbolic**: 0.12 ± 0.58°, *t*(70) = 0.20, *p* = 0.841, *d* = 0.02), suggesting that there was no implicit contributing to performance.

Unlike Experiment 2, in the analysis including all participants, the symbolic group exhibited a markedly lower asymptote compared to the sensory group. This is likely because there were significantly more non-learners in the symbolic group compared to the sensory group (ᵡ^2^ (1, 121) = 15.972, p < 0.001; **Sensory**: 16% non-learners; **Symbolic**: 54% non-learners). When we analyzed only the data from learners, the difference between the two groups was much more subtle. Here, the symbolic group showed a disadvantage only in the early phase of learning (Figure 3e, learners only: *t_score_* > 5.53, *p_perm_* < 0.05, *d* > 0.2), with the two groups reaching a similar late level of late adaptation (Figure 3f; paired t-test comparing late adaptation vs baseline, **Sensory**: 56.62 ± 1.19°, *t*(41) = 47.68, *p* < 0.001, *d* = 9.7; **Symbolic**: 56.47 ± 2.60°, *t*(32) = 21.22, *p* < 0.001, *d* = 5.28). Both groups exhibited a large drop in hand angle from late to aftereffect phases (Figure 3f: **Sensory**: –57.17 ± 1.17°, *t*(41) = –48.89, *p* < 0.001, *d* = –9.2; **Symbolic**: –56.25 ± 2.67°, *t*(32) = –20.77, *p* < 0.001, *d* = –5.24) and neither group exhibited an aftereffect (Figure 3f: **Sensory**: –0.55 ± 0.65°, *t*(41) = –0.85, *p* = 0.402, *d* = –0.2; **Symbolic**: 0.21± 0.61°, *t*(32) = 0.35, *p* = 0.731, *d* = 0.1).

The results of Experiment 3 reinforce the notion that strategy discovery is hindered by symbolic compared to sensory feedback. However, once discovered, the strategy can be applied with comparable efficiency for the two groups.

## Discussion

The term sensorimotor control captures the interwoven manner in which animals have evolved to navigate and manipulate their environments: Sensory signals provide information that is used to not only select a goal-relevant action but also ensure that the selected action is optimally executed. For learning, error information arising from sensory feedback ensures that the sensorimotor system can readily adapt to changes in the environment or bodily states. Motor adaptation, at least in humans, can also be driven by symbolic feedback in which the error information is conveyed in a more indirect, abstract manner. In three well-powered studies, we examined how implicit and explicit learning processes are modulated by symbolic feedback. Consistent with previous reports, we found that adaptation with symbolic feedback was dominated by explicit strategy use, with minimal evidence of implicit recalibration. Moreover, even when the symbolic feedback conveyed information content matched to that of sensory feedback, adaptation was markedly slower. We postulate that the abstract and indirect nature of symbolic feedback may impede learning by disrupting strategic reasoning and/or strategic refinement.

### Motor adaptation in response to symbolic feedback is dominated by an explicit re-aiming strategy

Building on measures that have been shown to be diagnostic of implicit and explicit processes in studies with sensory feedback, the results of Experiment 1 indicate that adaptation based on symbolic feedback is dominated by explicit strategy use. Signatures of strategy use include the scaling of participants’ performance and reaction times with the size of the perturbations, along with negligible residual aftereffects once the perturbation was removed. These findings are consistent with previous research indicating that adaptation from symbolic feedback arises from explicit strategy use (Butcher & Taylor, 2018).

However, there are some prior reports indicating that symbolic feedback can also elicit implicit recalibration (Izawa & Shadmehr, 2011; Larssen et al., 2022; van Mastrigt et al., 2023). There may be several reasons for this discrepancy. First, the aftereffects reported in these previous studies may not represent implicit recalibration but rather reflect some residual strategy use during the aftereffect phase (Morehead & de Xivry, 2021). This is likely to occur if participants are not explicitly told to “disengage from any strategy use and move directly to the target” – instructions we used in our studies.

Second, the way in which a symbolically cued perturbation is introduced may impact the extent of implicit recalibration. In our studies, a large perturbation was introduced in an abrupt manner: Feedback scores that had indicated a high degree of accuracy suddenly dropped with the introduction of the perturbation. In other studies, the perturbation has been introduced in a gradual manner (Therrien et al., 2018; Uehara et al., 2019); for example, the window within which a movement must end to earn a favorable binary outcome is gradually shifted in one direction. Under such conditions, participants will show an appropriate change in heading angle, and for relatively small shifts (up to 10-15°), show a small residual implicit aftereffect (1-3°) (van Mastrigt et al., 2023).

The mechanisms supporting implicit adaptation in response to symbolic feedback introduced in a gradual manner are unclear. On one hand, implicit learning might be driven by use-dependent learning, which refers to a movement bias toward frequently repeated movements (Diedrichsen et al., 2010; Huang et al., 2011; Marinovic et al., 2017; Tsay, Kim, Saxena, et al., 2022; Verstynen & Sabes, 2011). However, in a previous study, the generalization pattern to untrained targets after learning showed no evidence of attraction toward the repeated movement location, contradicting the use-dependent hypothesis (van Mastrigt et al., 2023). On the other hand, implicit learning could result from proprioceptive recalibration, where symbolic feedback induces a bias in perceived hand position in the direction opposite to the perturbation (Cressman & Henriques, 2011; Tsay, Kim, Haith, et al., 2022; Zhang et al., 2024). For instance, static reports of perceived hand position would be biased in the opposite direction of the perturbation, a prediction that needs to be examined in future studies. Regardless of the mechanism, it is notable that in all three of our experiments, explicit strategy use in response to symbolic feedback was sufficient to nullify the perturbation without the need for implicit learning.

### Symbolic feedback reduces explicit strategy use compared to sensory feedback

The results of Experiments 2 and 3 showed that learning was impeded by symbolic compared to sensory feedback. Under conditions in which the two forms of feedback conveyed similar magnitude and directional information, symbolic feedback led to fewer learners as well as a reduced rate of learning compared to sensory feedback. This learning disadvantage was evident in response to both a rotational perturbation and a mirror reversal perturbation.

These results contrast with those of Butcher and Taylor (2018) who failed to detect learning differences between sensory and symbolic feedback groups. The discrepancy may be due to differences in statistical power: Whereas Butcher and Taylor recruited 12-18 participants per group, our experiments included over 60 participants per group. The larger sample size in our study may have provided greater sensitivity to detect differences between sensory and symbolic feedback. Another possible account of the discrepancy relates to the timing of the feedback. We delayed the feedback by 800 ms, under the assumption that this would negate any contribution from implicit recalibration. In contrast, Butcher and Taylor (2018) provided immediate feedback. As such, one would expect that both implicit and explicit processes would be operative in their sensory feedback condition. While it was possible that the operation of multiple processes to be additive or at least sub-additive, the interaction of the processes might reduce the overall rate of learning in Butcher and Taylor’s sensory feedback condition, resulting in a null effect when compared to their symbolic condition in which learning was purely reliant on an explicit process.

Why might symbolic feedback impair learning compared to sensory feedback? Our current theoretical understanding of strategy use in motor learning tasks is quite limited. We have recently proposed a “3R” framework for strategy use consisting of three fundamental processes: Reasoning, the process of understanding action-outcome relationships; Refinement, the process of optimizing sensorimotor and cognitive parameters to achieve motor goals; and Retrieval, the process of inferring the context and recalling a control policy (Tsay et al., 2023). We postulate that the abstract and indirect nature of symbolic feedback may impede learning by affecting strategic reasoning and refinement. (Given that we didn’t manipulate the learning context, we aren’t in a position to assess the impact of symbolic feedback on retrieval.)

Sensory and symbolic feedback may elicit different forms of reasoning. In response to sensory feedback, participants may adopt “inference over hypotheses” (Rule et al., 2020; Tenenbaum et al., 2011). For example, with each movement, the participants might test whether the perturbation is a rotation or a reversal, using sensory feedback to rapidly update their beliefs about each hypothesis. In contrast, with symbolic feedback, where information is more abstract and indirect and the perturbation size and direction are more uncertain, participants may struggle to generate hypotheses about the perturbation. Instead, they may rely on a more heuristic, albeit slower form of reasoning, where successful actions are repeated and unsuccessful actions are avoided (Therrien et al., 2016; van Mastrigt et al., 2020). Future studies using periodic probes of generalization to novel target locations could arbitrate between these mechanisms: Learning via hypothesis testing should generalize to untrained target locations, whereas learning via trial-and-error would only generalize locally to trained target locations (McDougle & Taylor, 2019).

Alternatively, symbolic and sensory feedback may differ in terms of refinement. By this view, we assume that sensory and symbolic learners come to understand the nature of the perturbation at a similar rate. However, participants receiving sensory feedback, with direct visual input about the size and direction of the perturbation, may be quicker to convert this knowledge into an accurate motor plan. In contrast, symbolic feedback, which conveys size and direction in a more abstract and indirect manner, may make it harder for learners to translate this information into a successful control policy, thus slowing strategic refinement. This refinement notion suggests that the learning disadvantage in response to symbolic feedback may be mitigated by an extended familiarization block to better acquaint participants with the spatial information conveyed by the symbolic feedback.

## Supplemental Section

**Figure S1.**
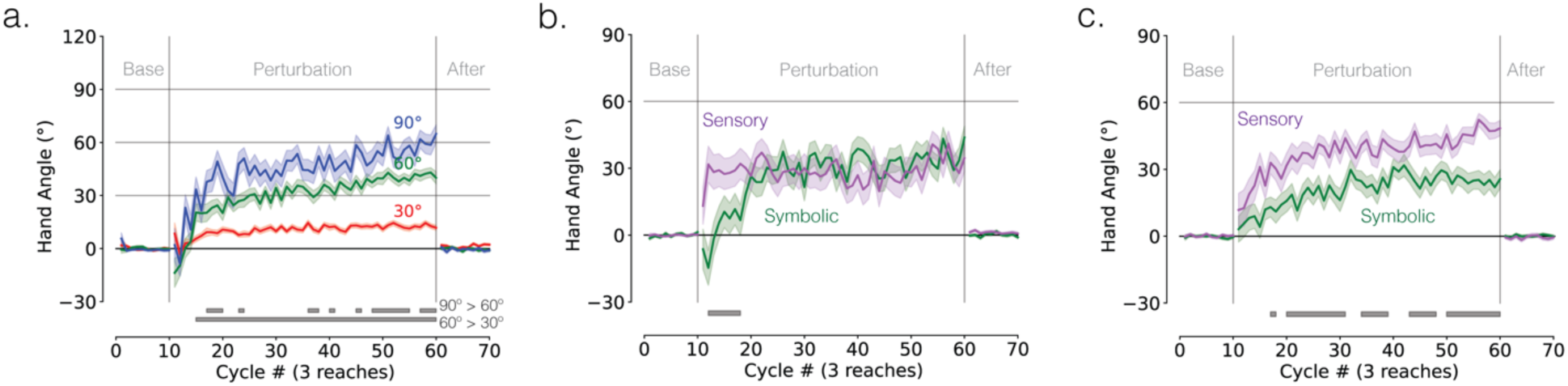
Right-handers only results. (**a**) Experiment 1: Mean time courses of hand angle (N = 156). Colors denote different perturbation groups (red = 30°; green = 60°; blue = 90°). Shaded error denoted SEM. Each movement cycle constitutes three reaching trials (1 reach to each of the three targets). Grey horizontal lines at the bottom indicate clusters showing significant group differences. **(b, c)** Mean time courses of hand angle for **(b)** Experiment 2 (N = 98) and **(c)** Experiment 3 (N = 106). Colors denote different feedback groups (dark green = symbolic feedback; dark magenta = sensory feedback).

**Figure S2.**
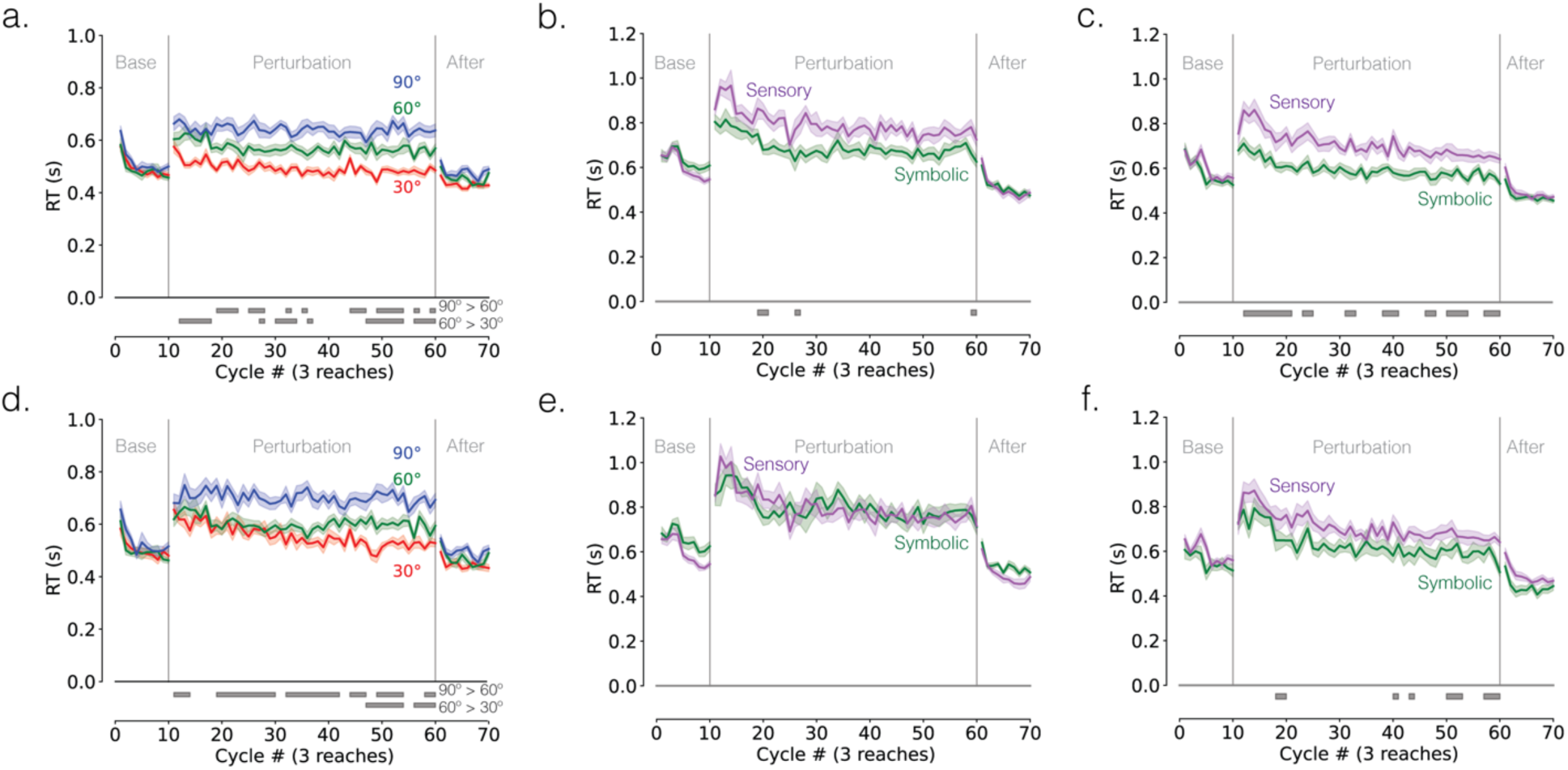
Reaction time (RT). (**a, d**) Experiment 1: Median time courses of RT (N = 184) for (**a**) all participants (N = 184) and (**d**) learners (N = 112). Colors denote different perturbation groups (red = 30°; green = 60°; blue = 90°). Shaded error denoted 95% Confident Interval. Each movement cycle constitutes three reaching trials (1 reach to each of the three targets). Grey horizontal lines at the bottom indicate clusters showing significant group differences. **(b, c, e, f)** Median time courses of RT for (**b**) all participants (N = 110) and (**e**) learners only in Experiment 2 (N = 78), (**c**) all participants (N = 121) and (**f**) learners only in Experiment 3 (N = 75). Colors denote different feedback groups (dark green = symbolic feedback; dark magenta = sensory feedback).

**Figure S3.**
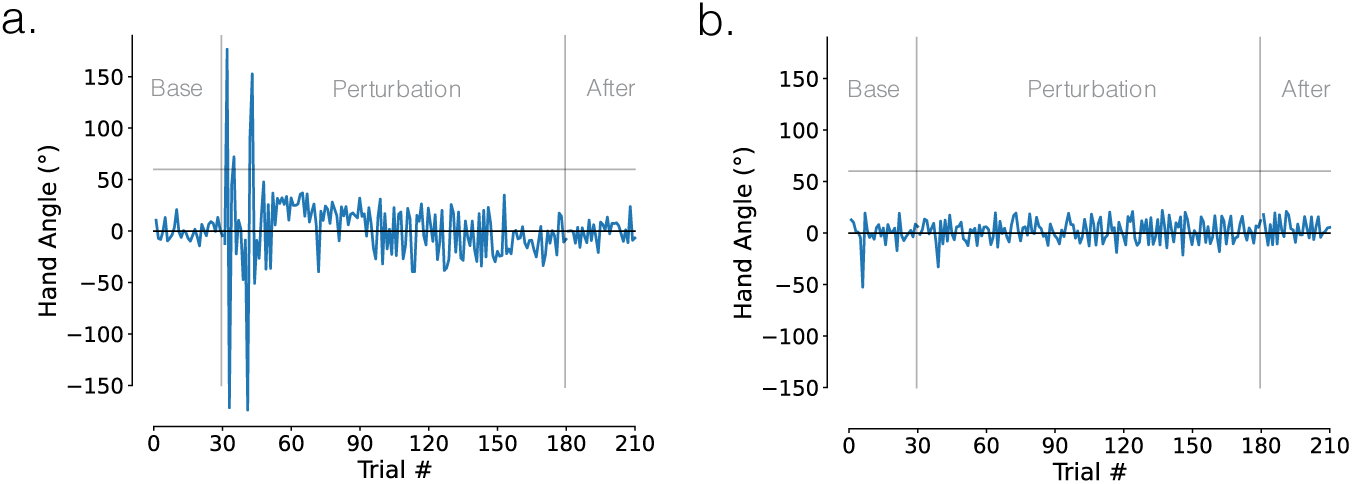
Representative non-learners in Experiment 1. (**a**) A representative non-learner who exhibited exploration early in learning but failed to identify a successful explicit strategy late in learning. **(b)** A representative individual who may have failed to follow task instructions, demonstrating little to no change in behavior during the perturbation block.

## Acknowledgements

We thank Jordan Taylor and Bev Larssen for their helpful comments on this manuscript. This project was supported by a Human Frontier Science Program post-doctoral fellowship awarded to JST and a NIH grant awarded to RBI (R35NS116883-01). The funders had no role in study design, data collection and analysis, decision to publish or preparation of the manuscript.

## Competing interests

RI is a co-founder with equity in Magnetic Tides, Inc., a biotechnology company created to develop a novel method of non-invasive brain stimulation. The other authors declare no competing interests.

## Notes

https://osf.io/bpfnh/?view_only=b1f4ba4e5576462c9be266380e2fee8b

### Competing Interest Statement

The authors have declared no competing interest.

